# Brassinosteroids function as the plant male and female reproductive hormone coordinating gene expression

**DOI:** 10.1101/2024.05.07.592278

**Authors:** Kumi Matsuura-Tokita, Takamasa Suzuki, Yusuke Kimata, Yumiko Takebayashi, Minako Ueda, Takeshi Nakano, Hitoshi Sakakibara, Akihiko Nakano, Tetsuya Higashiyama

**Author notes:** Corresponding author: Kumi Matsuura-Tokita. Author Contributions: K.M. and T.H. designed research; K.M., T.S., Y.T., H.S., performed Research; K.M., T.S., Y.K., Y.T., H.S., and M.U. analyzed data; and K.M., T.S., Y.K., M.U., T.N., H.S., A.N., and T.H. wrote the paper.

## Abstract

Brassinosteroids (BRs) are steroid hormones identified in plants. Besides promoting cell elongation and division, BRs facilitate the development of both male and female reproductive tissues. In animals, reproductive steroid hormones play an essential role in reproductive tissue development by regulating gene expression. Here, we focused on the function of BRs during fertilization. We measured the content of biologically active BRs, brassinolide (BL) and castasterone (CS), in the reproductive tissues of *Arabidopsis thaliana*. Both BL and CS accumulated abundantly in pollen grains and in larger amounts in pistils than in leaves. To evaluate BL function during fertilization, we used an *in vitro* guidance assay with exogenously applied BL. Although pollen tubes need to be elongated through the pistils for efficient capacitation, BL treatment promoted pollen tube capacitation and improved attraction to ovules *in vitro*. Transcriptome analysis demonstrated that BL treatment induced the expression of half of the genes expressed in pollen tubes that elongated through the pistils. These results indicated that BL supplied from pistils is a key factor for pollen tube capacitation. However, using the *bri1* mutant for the guidance assay resulted in reduced pollen tube capacitation, suggesting that BRI1-signaling in pistils is also important. Furthermore, BRs act on ovules. Exogenous BL application to ovules maintained guidance capacity by promoting the expression of small secreted proteins involved in pollen tube attraction and gamete fusion. Overall, BRs play a significant role as male and female reproductive hormones throughout the plant fertilization process.

## Introduction

Brassinolide (BL), the most active form of BRs, was originally isolated from the pollen grains of *Brassica napus L*. (1). BRs are enriched in the pollen and seeds of various angiosperm species and are known to promote cell elongation and division(2). In animals, steroid hormones such as androgens, estrogens, and progesterons play roles in reproductive organ development and stimulation. In insects, the steroid hormone ecdysone plays a role in ovarian development and oocyte maturation (3). The importance of BR signaling in anther/pollen and ovule development has been reported (4)(5), suggesting its functional similarity to animal steroid hormones. It has also been reported that pollen tube germination and growth are improved by exogenous BL application (6). In dicots, BL is produced from its precursor, castasterone (CS), by BR6OX2/CYP85A2, an enzyme that functions as a BR C-6 oxidase and BL synthase. CS is the second highest active form of BRs in dicots and is considered the final product in the BR biosynthesis pathway in monocots (7). Despite their implicated roles in reproduction, quantification of BRs in the reproductive tissues of a model plant, *Arabidopsis thaliana*, has not been conducted. Furthermore, the function of BRs during fertilization process in angiosperms remains unclear.

In angiosperms, communication between the male and female reproductive tissues is an essential cue for fertilization. Small peptides secreted from reproductive tissues play a significant role in this communication (8). For example, pollen tube germination in the stigma is regulated by RALFs secreted from the pollen tubes (RALF10/11/12/13/25/26/30) and pistils (RALF1/22/ 23/33) (9). Pollen tubes are capacitated only when they elongate through pistils, causing dramatic changes in gene expression (10)(11)(12)(13). RALF 4/19 peptides secreted from pollen tubes regulate growth by interacting with BUPS/ANX (14)(15). Once the pollen tube enters the ovules, RALF34 is secreted, which induces pollen tube rupture (16). Pollen tube attractants, LUREs, XIUQIUs, and TICKETs (17)(18)(19) are secreted from synergid cells in ovules. During the final stage of fertilization, thionins secreted from pollen tubes promote synergid cell degeneration (20)(11). EC1 peptides from egg cells promote gamete fusion (21)(22) and ECA1 gametogenesis-related family proteins (23) also play a role in fertilization. RALF6/7/16/36/37 secreted from pollen tubes (24) and ECS1/2 secreted from ovules (25)(26) are involved in blocking polytubey to accomplish fertilization. ECS1/2 have also recently been shown to be involved in synergid cell degeneration (26). However, the molecular mechanisms regulating these complex male-female communications remain to be elucidated.

In this study, we quantified the BL and CS content in *A. thaliana* reproductive tissues, pollen grains, and pistils. As previously reported, pollen grains contain high amounts of BL and CS. We found that the female reproductive tissue, the pistil, also contained BL and CS, although BL was not detectable in vegetative tissues (leaves). Physiological assays indicated that pollen tube capacitation and attraction by ovules were facilitated by exogenous BL addition. Transcriptome analysis revealed that gene expression profiles changed dramatically in both males and females, especially for small proteins involved in fertilization. Thus, BRs are reproductive hormones that regulate male-female communication during fertilization.

## Results

### BL and CS are highly concentrated in pollen grains

To identify the function of BRs during the reproductive process, we measured the content of BL and CS in male and female reproductive tissues (pollen grains and mature pistils) of *A. thaliana*. BL was highly concentrated in pollen grains, 109.9 +/- 13.2 pmol/gFW (Fig. 1A). CS content was also high at 84.3 +/- 13.3 pmol/gFW. These values are comparable to the BL content in pollen grains of *B. napus* L (1). However, in *B. napus* L., CS has not been detected. BL and CS in female tissues (pistils) were 0.47 +/- 0.15 and 3.33 +/- 0.58 pmol/gFW, respectively. BL was not detected in leaves. CS in leaves was 0.84 +/- 0.07 pmol/gFW.

**Figure 1.**
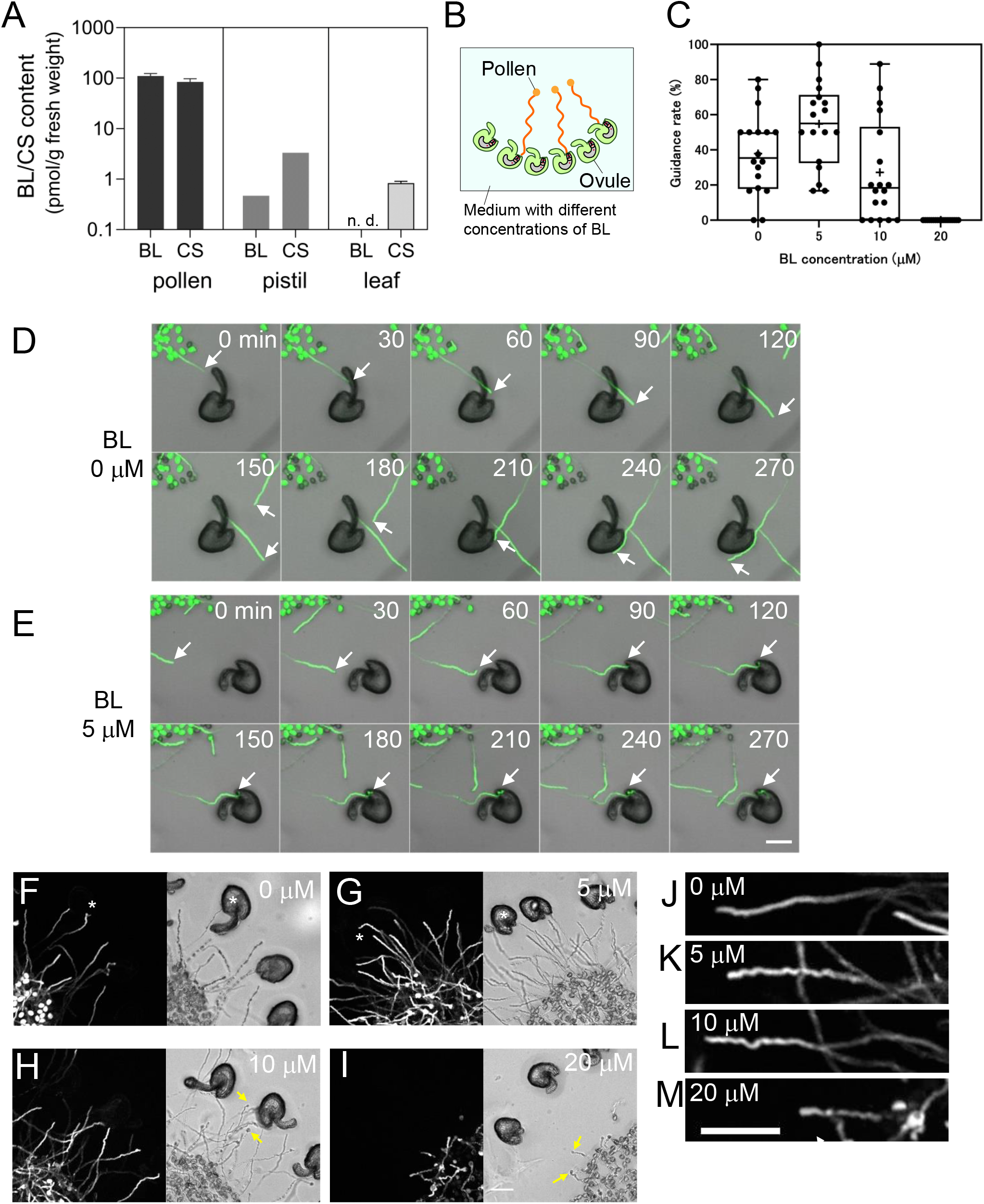
Effects of BL on pollen tube guidance. (A) Quantification of BL/CS in reproductive tissues of *Arabidopsis thaliana*. (B) Scheme of the *in vitro* guidance assay. (C) Quantification of guidance rate *in vitro* under different BL concentrations in pollen germination medium. With 5 μM BL, pollen tubes demonstrated the maximum guidance rate. (6 replicates, the total number of ovules was132 for each BL concentration). (D)Time-lapse observation of *in vitro* guidance assay with 0 μM BL. Pollen tubes approaching an ovule (white arrows) did not change the growth direction of the ovule. (E) Time-lapse observation of *in vitro* guidance assay with 5 μM BL. A pollen tube approaching an ovule (white arrows) changed the growth direction, entered ovule, and the tip burst. (F-I) Morphological differences of pollen tubes under different BL concentrations in PGM. With 5, 10, and 20 μM BL, pollen tubes demonstrated a wavy path. Pollen tube tips tended to burst with 10 and 20 μM BL (yellow arrows). Asterisks demonstrated ovules targeted by pollen tubes. (J-M) Enlarged view of pollen tubes in F-I with 0, 5, 10, and 20 μM BL. Scale bars, 100 μm.

### BL improved the capacitation of pollen tubes

Exogenous BL application promotes pollen tube germination and elongation *in vitro* (6). The optimal BL concentration for pollen tube growth was 10 μM, which is higher than that reported for hypocotyl elongation (0.1 μM) (27). The high optimal BL concentration for pollen tubes may reflect higher BL concentrations in pollen grains than in vegetative tissues (Fig. 1A). To test the function of BL during fertilization, we performed an *in vitro* quantitative assay for pollen tube guidance. When we exogenously applied BL to the pollen tube germination medium, the pollen tubes targeted the ovules more frequently. Surprisingly, pollen tubes were effectively activated without elongation through the pistils (Fig. 1 B, C). In the guidance assay, pollen grains expressing GFP in the cytosol were spread directly onto the agarose medium and incubated with the ovules. When pollen tubes entered an ovule and the pollen tube tip burst, the ovule was defined as “attracted.” Typical examples are shown in Fig. 1 D and E. With 0 μM BL, the pollen tubes tended to grow straight and pass through the ovule (Fig. 1D and Movie S1). With 5 μM BL, we observed that the pollen tubes changed their growth direction toward the ovule, entered the micropyle, and showed tip burst (Fig. 1E and Movie S2). The guidance rate was calculated as the percentage of ovules attracted. Pollen tubes treated with 5 μM BL demonstrated the highest guidance rate (Fig. 1C). However, when the BL concentration increased to > 10 μM, the guidance rate decreased. Thus, BL promoted pollen tube guidance to the ovules at an optimal concentration of 5 μM. BL treatment also affected pollen tube morphology. When BL was added to the pollen germination medium, pollen tubes grew in a wavy pattern in a dose-dependent manner (Fig. 1F-M). Such wavy pollen tubes have also been observed in capacitated conditions on the septum, or when growing toward the ovules. With BL concentrations more than 10 μM, pollen tube tip bursting was observed (Fig. 1H and I).

### BL treatment changed the expression profile of a wide variety of genes in pollen tubes

Recently, BL treatment has been shown to alter the expression profiles of various genes in roots (28). To explore whether the gene expression profile in pollen tubes was affected by BL treatment, we performed RNA-seq analysis. We compared three conditions: pollen tubes grown *in vitro* with BL (BL-PT), pollen tubes grown *in vitro* without BL (NC-PT), and pollen tubes elongated through the styles (SIV-PT) (Fig. 2, Dataset S1). In the BL-PT group, 263 genes were upregulated compared to those in NC-PT (log Fc > 1, FDR < 0.01, > 50 rpm). Among these, 238 genes (90.5 %) were also upregulated in SIV-PT compared to those in NC-PT. Also, 207 genes out of 445 genes (46.5 %) were upregulated only in SIV-PT (Fig. 2F). By contrast, approximately half of the genes (61.0%) downregulated with BL (logFC < −1, FDR < 0.01, > 50 rpm) showed overlap between BL-PT and SIV-PT (Fig. 2G). Gene Ontology (GO) enrichment analysis demonstrated that upregulated genes in Biological Processes (BP) were mainly involved in terms related to ion transport and pollen tube growth. Upregulated genes in the Cellular Component (CC) were highly involved in the term extracellular region (Fig. 2J). Characteristic genes (logFc > 0, FDR < 0.01, logCPM > 0.2) (Dataset S2) are listed in Table S1. These include CHX families and calcium-related genes. CNGC7, reported to be essential for male fertility, was upregulated (logFc = 0.91, FDR < 0.01) (29). In line with the improved guidance rate, PRK6, a LURE receptor, was also upregulated (logFc = 0.62, FDR < 0.01). Small peptides such as RALFs (RALF6/7/11/12/13/16/36/37), defensin-like proteins, and thionins were included. Other upregulated genes were related to cytoskeletal organization, cell wall, membrane traffic, and transcription factors like MYB120. Cell wall-related genes included callose synthases CALS9/10/12. CALS9/10 play roles on microspores to promote mitosis, and CALS12 is involved in microspore development (30). CALS10 was found to be enriched in sperm cells (31). CALS5, previously identified as responsible for callose plug formation in pollen tubes, (32) was not upregulated. However, we observed enhanced callose plug formation in BL-treated pollen tubes (Fig. S1). DMP9, a sperm cell-specific gene responsible for gamete fusion, was also upregulated (logFc = 0.54, FDR < 0.01) (33)(34)(35). By contrast, GO enrichment analysis indicated that downregulated genes in CC were involved in terms related to pollen tube tip growth (Fig. 2K).

**Figure 2.**
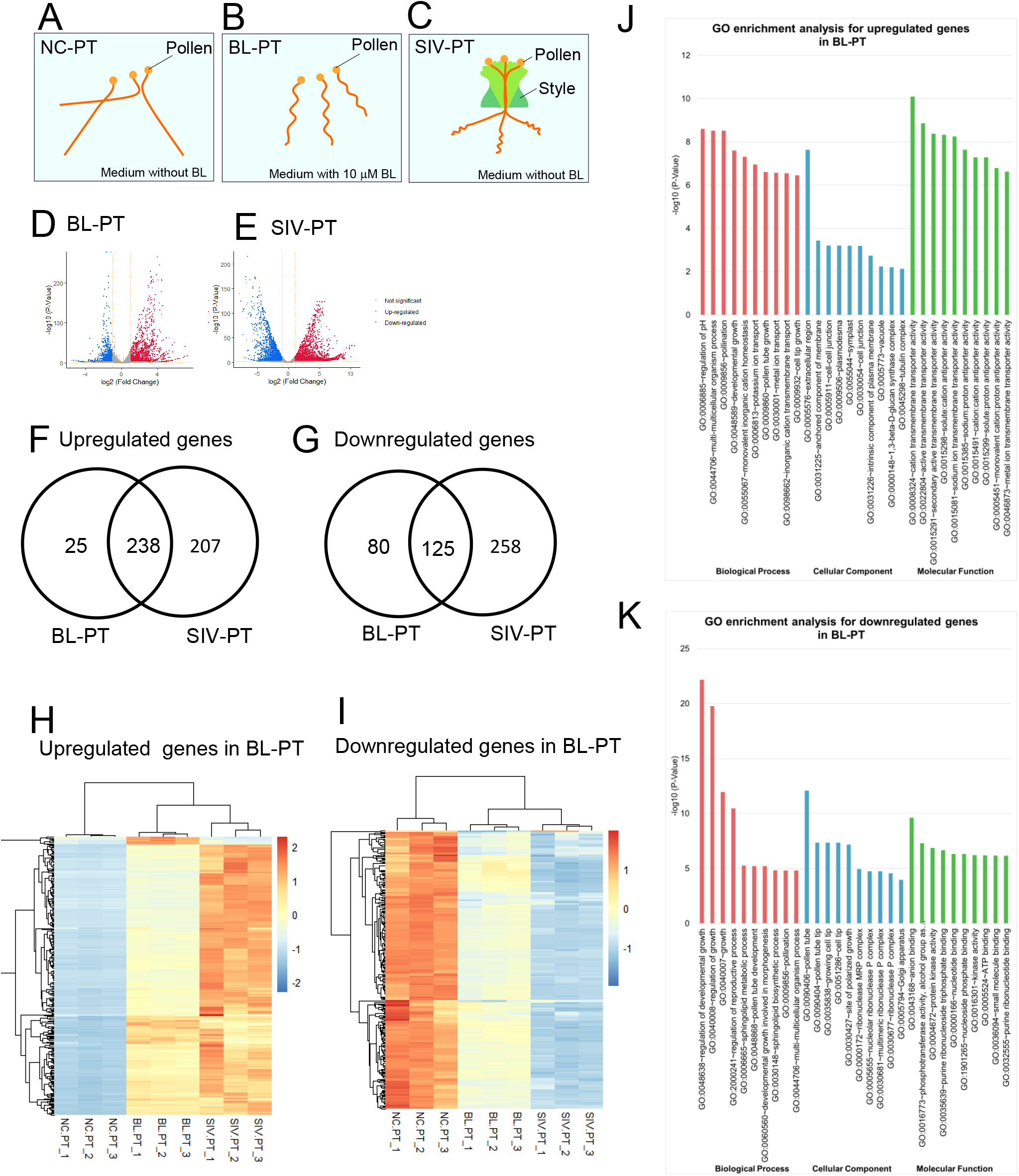
Transcriptome analysis of pollen tubes. (A-C) Schematic diagram showing different conditions for pollen tube incubation, (A) with 0 μM BL, (B) 10 μM BL, and (C) elongated through styles. (D) A volcano plot of differentially expressed genes in BL-PT compared with NC-PT. (E) A volcano plot of differentially expressed genes in SIV-PT compared with NC-PT. (F) A Venn diagram of upregulated genes (logFC > 1, FDR < 0.01, > 50 rpm) in BL-PT and SIV-PT. (G) A Venn diagram of downregulated genes (logFC < −1, FDR < 0.01, > 50 rpm) in BL-PT and SIV-PT. (H) A heatmap of upregulated genes in BL-PT. (I) A heatmap of downregulated genes in BL-PT. (J) GO enrichment analysis of upregulated genes in BL-PT using DAVID. (K) GO enrichment analysis of downregulated genes in BL-PT using DAVID.

Characteristic genes (logFc < 0, FDR < 0.01, logCPM > 0.2) (Dataset S3) are listed in Table S2. Downregulated genes included VGD1, PRK2/4/5, ANX2, BUPS1, and PICALM5a/b, all of which are implicated in pollen tube integrity and growth. GO enrichment analysis of genes upregulated only in the SIV-PT group revealed terms related to response to unfolded protein, cell junction, and clathrin-coated vesicles (Fig. S2) (Dataset S4). These data were in agreement with results of the physiological assay, which showed that pollen tubes were capacitated by BL-treatment (Fig. 1).

The transcriptome analysis also indicated that BL synthesis was promoted in BL-PT and SIV-PT (Fig. S3). We compared the expression of two genes involved in the final steps of BL synthesis, CYP85A1 and CYP85A2. Both genes are involved in castasterone (CS) synthesis. Of these, only CYP85A2 was found to mediate CS-to-BL synthesis. In growing pollen tubes, CYP85A1 expression was not detected in NC-PT and BL-PT, and it was scarcely detected in SIV-PT. In contrast, CYP85A2 expression was highly upregulated in BL-PT and SIV-PT and was extremely prominent in SIV-PT. The average expression of CYP85A2 in SIV-PT was more than ten times higher than that in NC-PT. These data suggest that BL is continuously synthesized in growing pollen tubes, especially in BL-PT and SIV-PT.

### BL pre-treatment of ovules maintained guidance capacity

In the results of the physiological assay (Fig. 1), both pollen tubes and ovules were treated with BL. To test whether exogenous BL application also affects female tissues, we conducted a guidance assay with ovules pre-treated with BL. Ovules were dissected from pistils and pre-incubated on agarose medium with or without 10 μM BL for 4 h. Ovules freshly dissected from the pistils were used as positive controls. After pre-treatment, the ovules were transferred to new agarose medium with BL and subjected to *in vitro* guidance assay. 10 μM BL-treated ovules maintained better guidance capacity than 0 μM BL-treated ovules (Fig. 3A, B).

**Figure 3.**
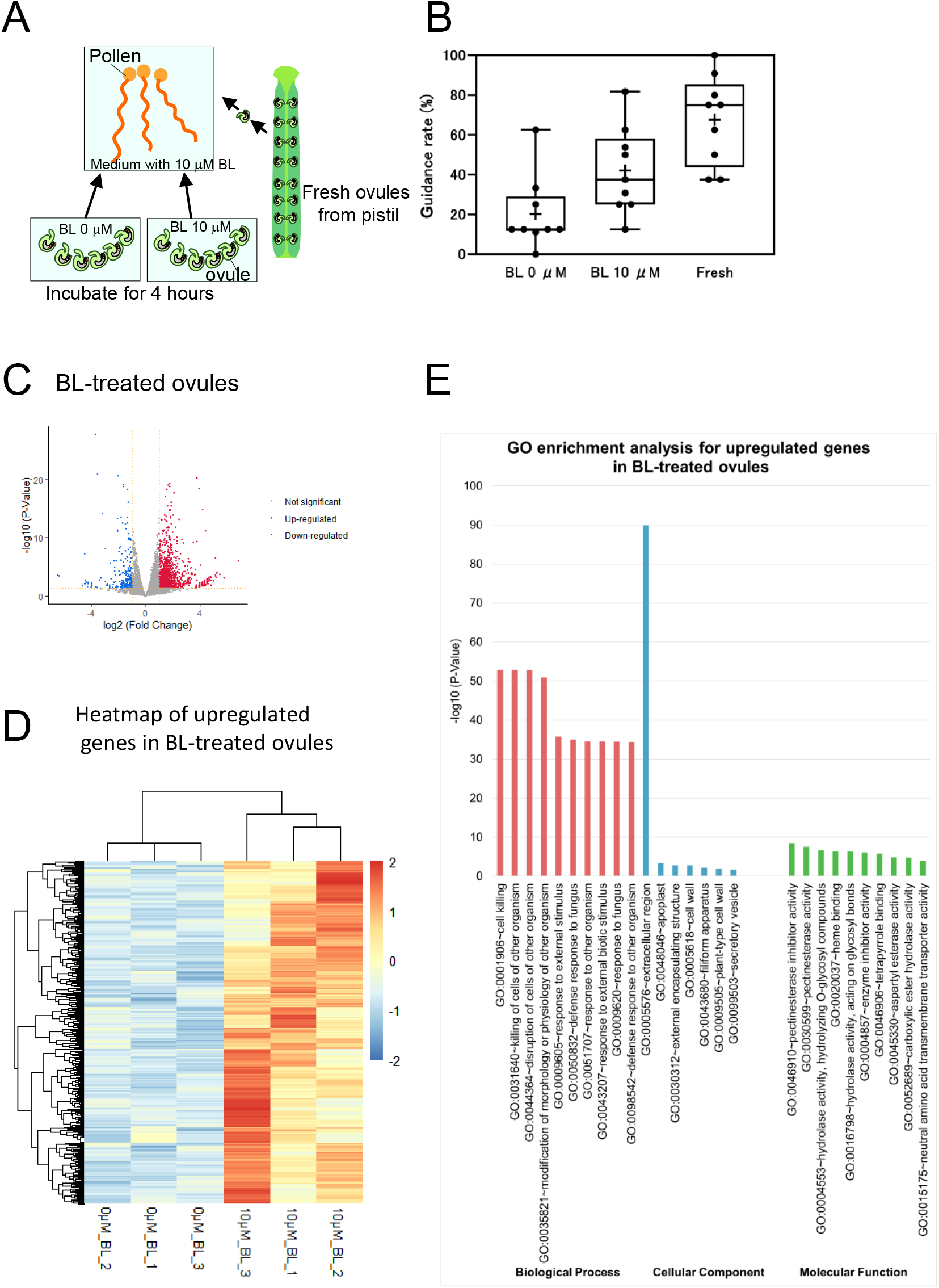
Effects of exogenously applied BL on ovules. (A) Scheme of the *in vitro* assay. Pollen tubes were incubated in PGM with 10 μM BL for 4 h before application of ovules. Ovules freshly dissected from pistils or incubated on medium with 0 or 10 μM BL for 4 h were applied to pre-incubated pollen tubes. (B) Quantification of guidance rates *in vitro* with ovules under different conditions. Freshly dissected ovules demonstrated the highest guidance rate, whereas ovules incubated with 0 μM BL lost guidance ability. Ovules incubated with 10 μM BL maintained guidance ability to some extent (3 replicates, the total numbers of ovules at each BL concentration were 79, 74, and 85, respectively). (C) A volcano plot of differentially expressed genes in BL-treated ovules. (D) A heatmap of upregulated genes in BL-treated ovules. (E) GO enrichment analysis of upregulated genes in BL-treated ovules using DAVID.

### BL pre-treatment of ovules induced gene expression of small proteins responsible for fertilization

Next, we performed RNA-seq analysis of ovules treated with BL for 4 h (Fig. 3C-E, Dataset S5). A total of 542 genes were upregulated in 10 μM BL-treated ovules compared to 0 μM BL-treated ovules (logFc >1, FDR < 0.05, > 1rpm). GO enrichment analysis indicated that the upregulated genes in BP were involved in terms related to defense responses. These upregulated genes in the Cellular Component (CC) were extensively involved in the term extracellular space.

Characteristic genes are listed in Table S3 (logFc > 0, FDR < 0.05, logCPM > 0.2) (Dataset S6). Small secreted proteins were abundant. These included pollen tube attractants such as LURE 1.2-1.8, XIUQIU 1-3, and XIUQIU4/TIC3. EC1 proteins involved in gamete fusion such as EC1.1, 1.3-1.5, ECA1 family proteins, defensin-like family proteins, and ECS1/2 were also upregulated. A comparison of our results with those of a previous single-cell RNA-seq analysis (36) revealed that genes specific to synergid cells, egg cells, and central cells were upregulated by BL-treatment (Fig. S4). Accordingly, the expression of MYB98, which is involved in the formation of the filiform apparatus from which pollen tube attractants are secreted (37), was upregulated. These data agreed with those from the physiological assay, which showed that BL pretreatment of ovules maintained guidance capacity (Fig.3 A, B).

### BRI1 signaling in pistils was important for pollen tube capacitation

Next, we performed semi-*in vivo* guidance assays using female reproductive tissues of mutants with aberrant BR signaling and synthesis. *bri1-116* is a mutant of the BR receptor BRI1. *cyp90a1-1* is defective in BL synthesis. *bil1-1D/bzr1-1D is a* dominant mutant in BRI1 signaling. To test the effects of BL on female tissues, we used a combination of wild-type and mutant ovules and/or styles (Fig. 4 A-D). When *bri1-116* or *cyp90a1-1* ovules were used, the pollen tube guidance rate decreased dramatically (Fig. 4B, C). By contrast, when using *bil1-1D/bzr1-1D* ovules, the guidance rate increased (Fig. 4D). Moreover, in the guidance assays using *bri1-116* ovules, we observed polytubey higher than that in wild-type (WT) ovules. Occasionally, inhibition of pollen tube entry at the micropyle was observed in *cyp90a1-1* and *bri1-116* ovules (Fig. S5). These results imply that both BR synthesis and the BRI1 signaling pathway are crucial for ovule activation.

**Figure 4.**
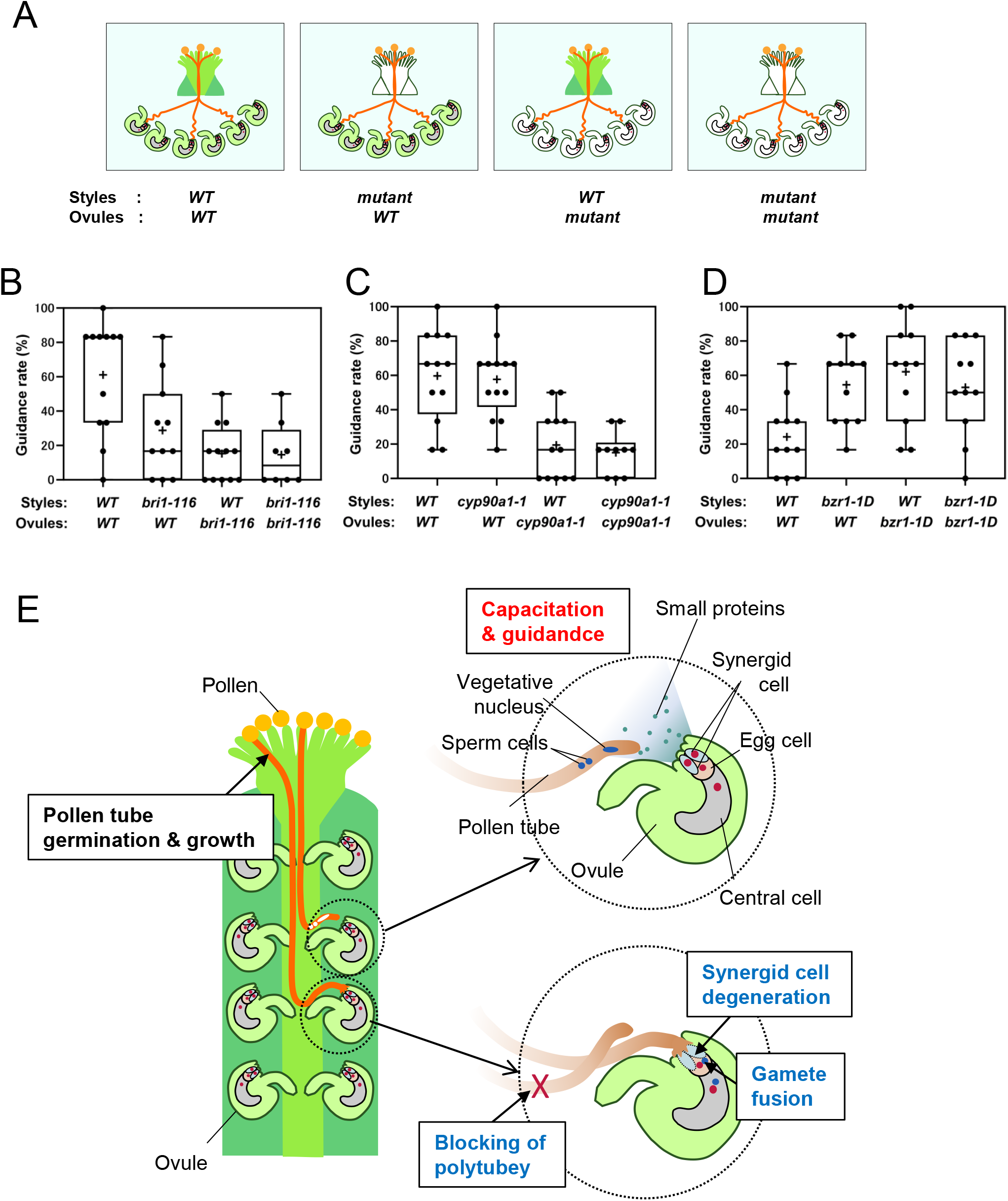
(A) Scheme of the semi-*in vivo* guidance assay using mutant styles and/or ovules. (B) Quantification of semi-*in vivo* guidance rate using *bri1-116* styles or ovules (4 replicates, the total number of ovules at each BL concentration were 72, 72, 66, and 48, respectively). (C) Quantification of semi-*in vivo* guidance rate using *cyp90a1-1* styles or ovules (4 replicates, the total numbers of ovules at each BL concentration were 72, 72, 78, and 60, respectively). (D) Quantification of semi-*in vivo* guidance rate using *bil1-1D/bzr1-1D* styles or ovules (4 replicates, the total numbers of ovules at each BL concentration were 66, 66, 66, and 66, respectively). (E) Roles of BR during fertilization. The red letters indicate BR functions revealed by physiological assay in this study. The blue letters indicate BR functions revealed by the transcriptome analysis in this study.

Interestingly, assays using the *cyp90a1-1* style demonstrated guidance rates comparable to those using the WT style, whereas the *bri1-116* style demonstrated a decreased guidance rate. By contrast, *bil1-1D/bzr1-1D* and WT ovules demonstrated increased guidance rates. A previous report showed that CYP90A1 promoter activity was detectable in stigma and transmitting tracts in pistils, and that BR was supposed to be synthesized with pollen tube elongation (6). In conclusion, these results suggest that BRI1 signaling in pistils, not BR synthesis, is important for pollen tube capacitation *in vivo*.

### BRI1 signaling is important for filiform apparatus development in ovules

A recent study showed that approximately half of *bri1* mutants were defective in embryo sac development (5).To investigate the relationship between structural defects and guidance ability, we performed electron microscopic observations of *bri1* mutant ovules. I In *bri1-116 brl1 brl3* wox5-GFP (*bri1*-triple) mutant ovules, the region of the filiform apparatus (FA) in the synergid cells was smaller than that in the WT ovules (Fig. S6A-C). As FA is considered a secretion site for pollen tube attractants (38), abnormal FA development in the *bri1*-triple mutant might be a cause of pollen tube guidance defects. By contrast, ovules in the *bil1-1D/bzr1-1* mutant demonstrated protrusions in the FA region. Taken together, these data suggest that BRI1 downstream signals play a significant role in the secretion of small peptides for fertilization and ovule development.

## Discussion

In this study, we found that BL played significant roles throughout fertilization by regulating gene expression in male and female reproductive tissues (Fig 4E).

### BL/CS showed quantitative differences in reproductive tissues

We measured the BL and CS content in the reproductive tissues of *A. thaliana*. In accordance with previous reports, BL and CS were highly concentrated in pollen grains. In *B. napus L*., only BL was found in pollen. However, in *A. thaliana*, we found almost the same amounts of both BL and CS in pollen. The expression of CYP85A2, an enzyme responsible for BL synthesis from CS, was more than ten times higher in SIV-PT than in NC-PT. As upregulation of CYP85A2 was not so prominent in BL-PT, elongation of pollen tubes through pistils was crucial for high level of CYP85A2 expression.

We detected both BL and CS in larger amounts in pistils than in leaves. These BRs may function in pollen tube capacitation and small protein secretion during fertilization.

### BL capacitated pollen tubes *in vitro*

The capacitation of PTs is a key process in angiosperm reproduction. In this and previous studies(39)(40)(41), *in vitro* germinated pollen tubes (pollen tubes germinated on agarose medium) were not very sensitive to ovular guidance signals. Contact with female tissues (pistils) dramatically changes the gene expression patterns of pollen tubes (42)(11)(10). However, our physiological assays and transcriptome data showed that BL can capacitate pollen tubes without contact with female tissues. The main factors involved in pollen tube capacitation might be members of the CHX family and calcium-related factors involved in ion transport and signal transduction. Factors related to the cytoskeleton, cell wall, and membrane traffic also demonstrated changes in expression. Exogenous application of BL promoted callose plug formation in pollen tubes (Fig. S1). As previous study reported that mutants lacking callose plugs showed reduced fertility when competed with wild type pollen tubes (32), there is a possibility that enhanced callose plug formation contribute to improved fertilization. BL promoted expression of CALS9/10/12, which play roles in pollen development and mitotic stage. As phylogenetic analysis of callose synthase revealed that CALS9/10 and CALS11/12 diverged after the divergence of angiosperms (31), these CALS may have specific roles in angiosperm-specific fertilization processes.

Consistent with the increased pollen tube guidance rate, the expression of PRK6, a receptor for pollen tube attractants, was also upregulated. However, approximately half of the genes upregulated in SIV-PT were not upregulated in BL-PT, suggesting the existence of other physiologically active substances in the pistils that capacitate pollen tubes. Interestingly, downregulated genes in BL-PT (Table S2) contained previously identified genes involved in pollen tube integrity and growth, such as VGD1, PRK2/4/5, ANX2, BUPS1 and PICALM5a/b (43)(44)(45)(46)(47). BL treatment may change the condition of pollen tubes from a “growing state” to a “fertilization state”. Previously identified in pollen tube capacitance in *Torenia*, AMOR is an arabinogalactan sugar chain secreted from ovules (48). To date, AMOR has not been identified in *A. thaliana*; however, AGPs and glycoprotein families with arabinogalactan sugar chains are involved in reproduction (49)(50). AGPs are expressed in pistils along the path of pollen tubes and are expected to play a role in pollen-pistil interactions (51). We also detected elevated expression of AGP5/15/23 in ovules after BL treatment. Further studies are required to explore their roles in pollen tube capacitation.

### BL facilitates communication between males and females

BL treatment of ovules promoted the expression of various small secreted protein genes (Fig. 3). These include pollen tube attractants from synergid cells, EC1 from egg cells, and ECA1 family proteins from synergid and central cells. Electron microscopic observation of ovules in mutants related to BRI1-signaling (Fig. S6) demonstrated that formation of the filiform apparatus is affected by reception of BRs. Transcriptome analysis demonstrated that MYB98 expression was BL-dependent (Table S3). BL and CS contained in the pistils may contribute to these developmental processes. BL treatment also promoted the expression of small proteins such as RALFs, defensin-like proteins, and thionins in pollen tubes. Thionins were reported to be downstream factors of MYB120 (11). Recent studies have demonstrated the importance of RALF peptides in fertilization(24)(9). Among them, RALF11/12/13, involved in interspecific barriers, and RALF6/7/16/36/37 which prevent polytubey, were BL-dependent. EC1 facilitates gamete fusion by relocating HAP2/GCS1 to the sperm cell surface. Interestingly, BL-treatment upregulates DMP9, a sperm cell-specific gene involved in gamete fusion, together with EC1 and HAP2/GCS1 (33)(34)(35). Physiological analysis using *bri1*mutant styles showed BRI1 signaling in pistils is important for pollen tube capacitation. Assays using *bri1* and *cyp90a1* mutant ovules sometimes demonstrated polytubey and inhibition of pollen tube entry at the micropyle. Elucidating whether these phenomena arise from structural defects in mutant ovules or defects in male-female communication will be an intriguing pursuit. In conclusion, we showed that expression of various genes involved in fertilization was controlled by a steroid hormone, BR. Dissecting distinct BR signaling pathways in male and female tissues is the next issue for research.

### Materials and Methods Strains

*A. thaliana* ecotype Columbia (*Col-0*) was used as the WT. *cyp90a1-1* (SALK_023532), *bri1-116, bri1-10* (SALK_041648), *bri1-triple* (*bri1-116brl1brl3 wox5-GFP)*, and *bil1-1D/bzr1-1D* (52) were also used in this study. *cyp90a1-1* and *bri1-10* (SALK_041648) were purchased from Abcam (Cambridge, ABRC). *bri1-trple* (*bri1-116brl1brl3 wox5-GFP*) was gifted by Dr. Russinova. *bri1-116* and *bil1-1D/bzr1-1D* were gifted by Dr. T. Nakano. Plants were grown under 16 h light/8 h dark conditions. Generally, seeds were planted directly in the soil. In the case of T-DNA insertion lines from ABRC, seeds were sterilized with 97% ethanol and planted on 0.5× Murashige and Skoog medium (Duchefa Biochemie B. V.) containing 1% sucrose and 1% agar. After 2 d of growth in the dark at 4 °C, plates were incubated under continuous light at 22 °C for approximately 10 d and subjected to genotyping analysis, and then homozygous mutants were inoculated onto soil.

### Quantification of BL/CS

Pollen grains were collected using vacuum cleaner as described (53). Pollen grains in 5.3, 8.4, and 8.5 mg weight was used for quantification (3 replicates). For preparation of pistil samples, stage 12 flowers were emasculated one day before. Mature pistils from 50-100 flowers (9.1 and 9.8 mg) were collected (2 replicates). Leaves were collected and cut into small pieces. The samples weighed 97.5, 103.9. 91.0 mg, respectively (3 replicates). Subsequently, BL and CS were extracted and semi-purified as described previously (54) except that the frozen tissues were lyophilized and crushed to a fine powder with 5-mm zirconia beads by TissueLyser (Qiagen, Hilden, Germany). Quantification was performed according to the previous report with UHPLC-Q-Exactive (Thermo Fisher Scientific) (54) except for the use of an ODS column (AQUITY PREMIER HSS T3, 1.8 µm, VanGuard TM FIT, 2.1 x 100 mm; Waters) according to the following profile: 0 min, 90% A + 10% B; 2 min, 60% A + 40% B; 17 min, 35% A + 65%B; 17.1 min, 2% A + 98% B. Data were processed by Xcalibur TM 4.2.47 (Thermo Fisher Scientific).

### Pollen tube guidance assay

The pollen tube guidance assay was performed as described in Hamamura *et al*.(55). Pollen germination medium (PGM) was prepared as follows, 1% low-melting agarose (Lonza) medium containing 0.01% H_3_BO_3_, 1 or 5 mM CaCl_2_, 5 mM KCl, and 1 mM MgSO_4_ (56) was melted at 65°C, poured into wells of a 1 mm-thick silicone sheet, and allowed to settle for a few minutes to solidify. Epibrassinolide (Sigma-Aldrich E1641) dissolved in 90% ethanol was added to PGM when necessary (6). Stage 12 flowers were emasculated one day before pollination. Pollen grains or pollinated styles from stage 13 flowers were placed on PGM and incubated in a moist chamber at room temperature for 4–5 h. Pollen tubes were imaged using a CLSM (Olympus IX-81 inverted microscope equipped with a spinning disk confocal unit (CSU-X1; Yokogawa Electric), LD lasers (445 nm), and an EM-CCD camera (evolve512; Teledyne Photometrics)) using a 10× objective lens with Metamorph software (Molecular Devices). To visualize the behavior of pollen tubes, WT strains harboring LAT52p-sGFP (57) were used to visualize pollen tubes behaviors. Images were acquired every 10 min. All image analysis was performed using ImageJ and Fiji software (58)(58). For assays using *bil1-1D/bzr1-1D* styles or ovules, less pollen was used because the guidance rate was higher than that of WT.

### RNAseq sample preparation

For *in vitro*-grown pollen tube samples, pollen grains from freshly opened flowers (50 flowers per sample × 3 replicates) were spread on cellulose cellophane (#300 Futamura Chemical) on PGM. Pollen tubes were germinated and grown in a moist chamber at room temperature for 5 h. Cellophane sheets containing pollen tubes were collected and immediately frozen in liquid nitrogen. For semi-*in vivo* samples, stage 12 flowers were emasculated 24 h before the experiment, and 30 - 40 pistils per sample (3 replicates each) were excised, pollinated with stage 13 flowers, and incubated in a moist chamber for 5 h. PTs growing from sections of the styles were cut with 27 gauge × 3/4” needles (Terumo Corporation), collected in a tube containing lysis buffer (Dynabeads mRNA DIRECT Micro kit; Thermo Fisher Scientific K.K.), and immediately frozen. For ovule samples, flowers were emasculated as described above, 24 h before sampling. Ovules were collected from 2 to 3 pistils.

mRNA was extracted using a Dynabeads mRNA DIRECT Micro Kit. Library construction was performed using the TruSeq RNA Sample Preparation Kit ver. 2 (Illumina). Sequencing was performed using NextSeq500 (Illumina) with a NextSeq500 High Output Kit v2.5 (Illumina).

### Analysis of RNAseq data

Raw reads containing adapter sequences were trimmed using bcl2fastq (Illumina), and nucleotides with low quality (QV < 25) were masked by N using the original script. Reads of <50 bp were discarded, and the remaining reads were mapped to the cDNA reference sequence using Bowtie with the following parameters: ‘--all –best --strata’ (59). Reads were then counted by transcript models. RNA-seq data were analyzed using the R 4.0.2 statistical software.

Differentially expressed genes were identified using edgeR software (60). For pollen tubes, the fold-change threshold was log_2_Fc >1 or <-1, and the false discovery rate threshold was FDR <0.01. For ovules, the fold-change threshold was log_2_Fc >1 or <-1, and the false discovery rate threshold was FDR <0.05. GO analysis was performed by The Database for Annotation, Visualization and Integrated Discovery (DAVID) 2021 (61)(62).

### Transmission electron microscopy (TEM) of ovules

Ovules of the WT, *bri1-triple* (*bri1-116brl1brl3 wox5-GFP*), and *bil1-1D/bzr1-1D* strains were subjected to TEM analysis. Stage 12 flowers were emasculated 24 h before fixation. Pistils were dissected using insect pins, and the ovary walls were removed to isolate the ovules. Isolated ovules were fixed with 2% paraformaldehyde (PFA) and 2% glutaraldehyde (GA) in 0.05M cacodylate buffer (pH 7.4) overnight at 4 °C. After fixation, ovules were washed three times with 0.05 M cacodylate buffer for 30 min each and postfixed with 2% OsO_4_ in 0.05 M cacodylate buffer for 3 h at 4 °C. Ovules were dehydrated in ethanol solutions (50%, 70%, 90%, and 100%). The ovules were then infiltrated with propylene oxide (PO) and embedded in resin (Quetol-651; Nisshin EM Co.). Ultrathin sections (80 nm) were stained with 2% uranyl acetate followed by a lead staining solution (Sigma-Aldrich Co.) and observed using a transmission electron microscope (JEM-1400Plus; JEOL Ltd.). Digital images were captured using a CCD camera (EM-14830RUBY2, JEOL Ltd.). Five ovules were observed for each strain,

## Acknowledgments

We would like to thank Yoko Mizuta for technical support and for providing the wild-type strains harboring LAT52p-sGFP; Daisuke Kurihara, Yoshikatsu Sato, Masahiro Kanaoka, Narie Sasaki, Akane Mizukami, and Satohiro Okuda for technical support; Mikiko Kojima and Ayami Furuta for technical assistance; Eugenia Russinova for providing the *bri1-triple* (*bri1-116brl1brl3 wox5-GFP*) mutant; and Editage (www.editage.jp) for English language editing.

This work was supported by the Japan Society for the Promotion of Science (JP15J40125 to K.M., JP18K19333 to T.H., JP21K18235 to T.H. and K.M., 22H04980 to T.H., and 22K21352 to T.H.). and CREST, Japan Science and Technology Agency (JPMJCR20E5 to T.H.).

